# Hebbian activity-dependent plasticity in white matter

**DOI:** 10.1101/2022.01.11.473633

**Authors:** Alberto Lazari, Piergiorgio Salvan, Michiel Cottaar, Daniel Papp, Matthew Rushworth, Heidi Johansen-Berg

**Affiliations:** Wellcome Centre for Integrative Neuroimaging, FMRIB, Nuffield Department of Clinical Neurosciences, University of Oxford; Wellcome Centre for Integrative Neuroimaging, Department of Experimental Psychology, University of Oxford

## Abstract

Synaptic plasticity is required for learning and follows Hebb’s Rule, the computational principle underpinning associative learning. In recent years, a complementary type of brain plasticity has been identified in myelinated axons, which make up the majority of brain’s white matter. Like synaptic plasticity, myelin plasticity is required for learning, but it is unclear whether it is Hebbian or whether it follows different rules. Here, we provide evidence that white matter plasticity operates following Hebb’s Rule in humans. Across two experiments, we find that co-stimulating cortical areas to induce Hebbian plasticity leads to relative increases in cortical excitability and associated increases in a myelin marker within the stimulated fiber bundle. We conclude that Hebbian plasticity extends beyond synaptic changes, and can be observed in human white matter fibers.

## Introduction

Hebb’s Rule [1] has been extremely influential in neuroscience. It postulated for the first time that a computational principle could link a biological process (“neurons that fire together, wire together”) with a cognitive process (Pavlovian/associative learning), an idea that has become pivotal for neuroscience research. Hebb’s Rule was later found to have a biological substrate in the synapse. Synapses can detect coincident activity of two neurons, i.e. detect when neurons “fire together”, and effect plastic changes in the synaptic connections between them, i.e. make neurons “wire together” [2]. Strikingly, more than half a century after it was first proposed, Hebbian theory is still thought to be accurate, although it is now encompassed by wider frameworks such as Spike-Timing Dependent Plasticity [3] or Bienenstock-Cooper-Munro theory [4]. In addition, extensive evidence has demonstrated that synaptic plasticity and its Hebbian properties are crucial for learning [5,6].

In recent years, another key site of brain plasticity has been identified: the myelinated axon [7]. Myelinated axons make up the majority of brain’s white matter, where this form of plasticity was first identified in humans [8]. This distinct plastic process has been confirmed to have two properties similar to synaptic plasticity: it is activity-dependent and it is implicated in learning. Its activity-dependence has now been confirmed in animal models across a broad range of methods, including electrical stimulation [9], optogenetics [10], chemogenetics [11], prevention of synaptic vesicle release by tetanus toxin [12] and non-invasive Transcranial Magnetic Stimulation [13]. Regarding its link to behaviour, active myelination is critical for a wide range of learning behaviours [14], including motor learning [15], fear learning [16], and spatial memory [17].

However, unlike synaptic plasticity, myelin plasticity has not been directly linked to known computational principles, and it is still unclear what rules it might follow. Our study was designed to test whether myelin plasticity follows Hebb’s Rule. To induce short-term plasticity, we used non-invasive Transcranial Magnetic Stimulation (TMS) to elicit neuronal activity with tight temporal control over two brain areas in a Hebbian fashion [18, 19]. We then combined Hebbian stimulation with MR-based quantitative myelin markers to detect myelin changes induced by Hebbian stimulation.

To facilitate a biological interpretation of our results, we focussed on Magnetisation Transfer saturation (MT), an MR-based metric that has been extensively validated with myelin histology [20, 21]. Unlike in rodent experiments, in which it is possible to carefully control the environments of experimental subjects, the possibility of variation in environment across human participants meant that care had to be taken to ensure that measures of physiology and myelination would have the best chance of revealing any impact of plasticity that might have occurred. We therefore scanned participants 24 hours before and after Hebbian stimulation. This time frame was selected to be sensitive to both the physiological changes that are associated with Hebbian plasticity (which are apparent soon after Hebbian stimulation [18]), and myelination-related effects, such as remodelling of myelin morphology, changes in the length of Nodes of Ranvier, or production of myelin from existing oligodendrocytes [22, 23, 24, 25]. These myelination-related effects may take slightly longer but can occur within 24 hours [7, 22, 23, 24, 25], and are all known to impact MT measurements [20, 21].

## Results

### Inducing Hebbian plasticity in the human brain

We used Hebbian stimulation in healthy adult participants to induce associative plasticity between right ventral premotor cortex (PMv) and left primary motor cortex (M1) (Fig.1A-B, Fig.S1, Fig.S2; see also Methods, Experimental design). First (study 1: Hebbian; fig.1A, B), we checked that a protocol which has been shown to induce Hebbian plasticity [18, 19] induced a measurable change in the excitability of M1 between the two testing days; this was indeed the case (left bar, figure 1C; subset of n=7 participants from Study 1). We then repeated the same testing protocol in a separate group of 18 participants (study 2: Hebbian) and compared it with a control procedure, which we refer to as “non-Hebbian stimulation”, conducted in another group of 18 participants (Study 2: non-Hebbian). When we compared changes in excitability in left M1 between the two testing days in the two conditions in study 2, we found there was a clear difference (Fig. 1C, Study 2:Hebbian vs Study 2:non-Hebbian, Mann-Whitney U test, p = 0.0049). While reductions in excitability over time were observed for the control condition (as expected from similar longitudinal studies [26]), this effect was rescued by the Hebbian plasticity-induction protocol, resulting in relatively greater M1 excitability following the Hebbian procedures. We then pooled together data from all participants, and found again that changes in cortical excitability of left M1 differed between the stimulation conditions (oneway ANOVA, F(2, 42) =8.747, p = 0.0126). This effect was present 24 hours after stimulation, indicating a long-lasting physiological effect of the stimulation compatible with the longer timescales expected in myelin plasticity [7].

**Figure 1:**
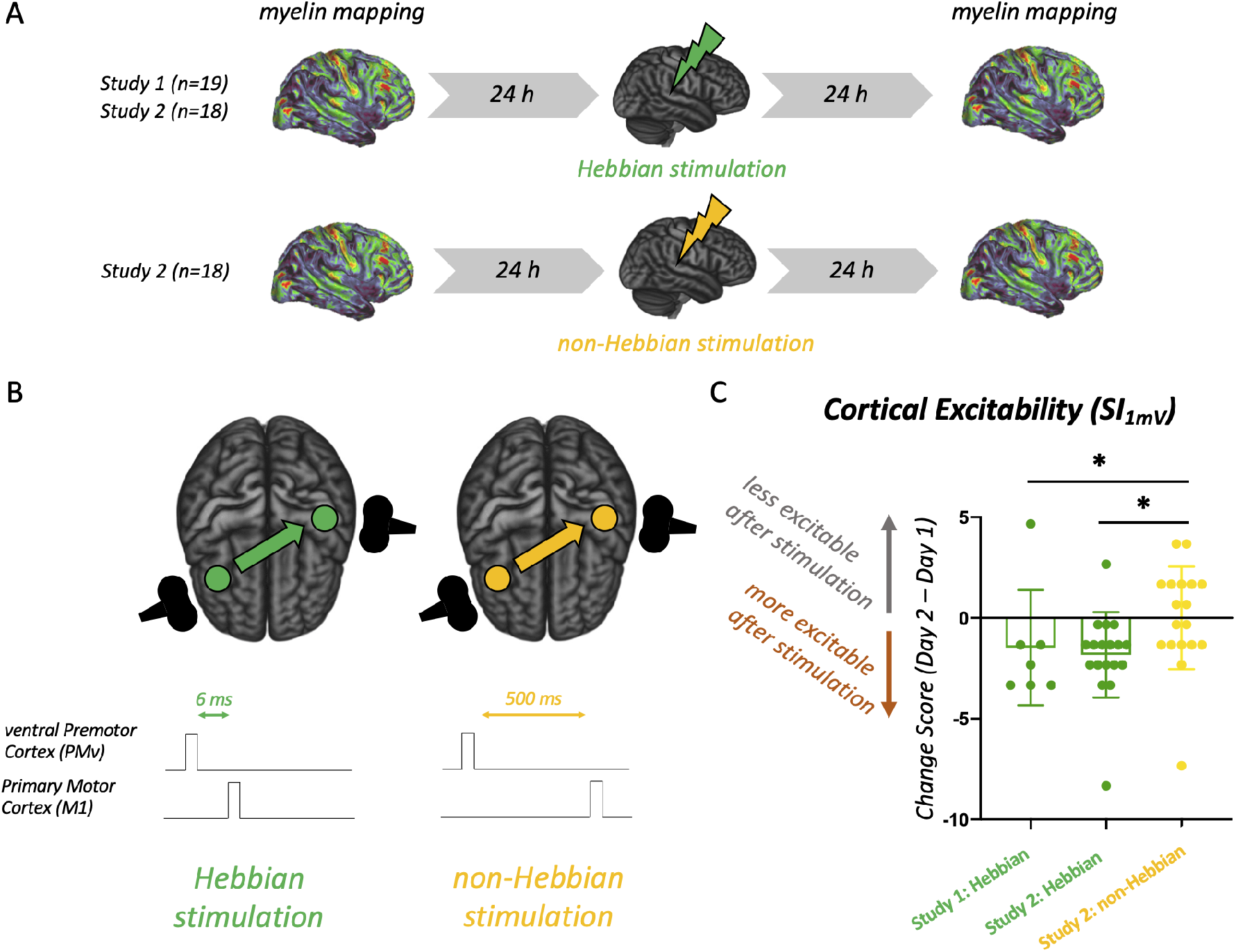
Inducing Hebbian plasticity in the human brain. A: Summary of experimental design, using two cohorts to establish effects of Hebbian stimulation on brain microstructure. Study 1 (n=19) included the Hebbian condition only. In Study 2, a different set of individuals were randomised to receive either Hebbian (n=18) or non-Hebbian (n=18) stimulation. *B:* Diagram of the Hebbian (active) and non-Hebbian (control) conditions used in the experiments. Both stimulation protocols are matched for duration, intensity and coil location, but differ in the relative timing of the stimulation pulses, with the Hebbian condition aiming to mimic the timing of synaptic plasticity inductions used *in vitro. C:* Longitudinal effects of Hebbian plasticity induction on cortical physiology. Each dot in the graph represents the normalized change in cortical excitability (as measured by the SI_1mV_ metric) for one subject. The SI_1mV_ measure was collected in an exploratory manner in the last 7 participants of study 1, and in all participants of study 2 to confirm the presence of longitudinal effects.

### Hebbian activity-dependent plasticity in white matter

To test whether Hebbian stimulation induced myelin plasticity, we collected highly reliable (Fig.S3) whole-brain myelin-sensitive Magnetisation Transfer saturation (MT) maps 24 hours before and after Hebbian stimulation in Study 1 and Study 2. Group comparisons of changes in MT did not detect significant differences between Hebbian and non-Hebbian conditions. However, using a whole-brain analysis, we were able to test whether physiological changes induced through Hebbian stimulation were associated with changes in myelin maps anywhere in the brain. We found a significant cluster in which participants with the strongest increases in cortical excitability, specifically following Hebbian stimulation, also exhibited the strongest increases in MT (Fig. 2A, peak p_corr_ = 0.013). In both studies, this effect was present in those receiving Hebbian stimulation (Fig. S4), but was not present in those receiving non-Hebbian stimulation (Fig.2C, Fig. 2E).

**Figure 2:**
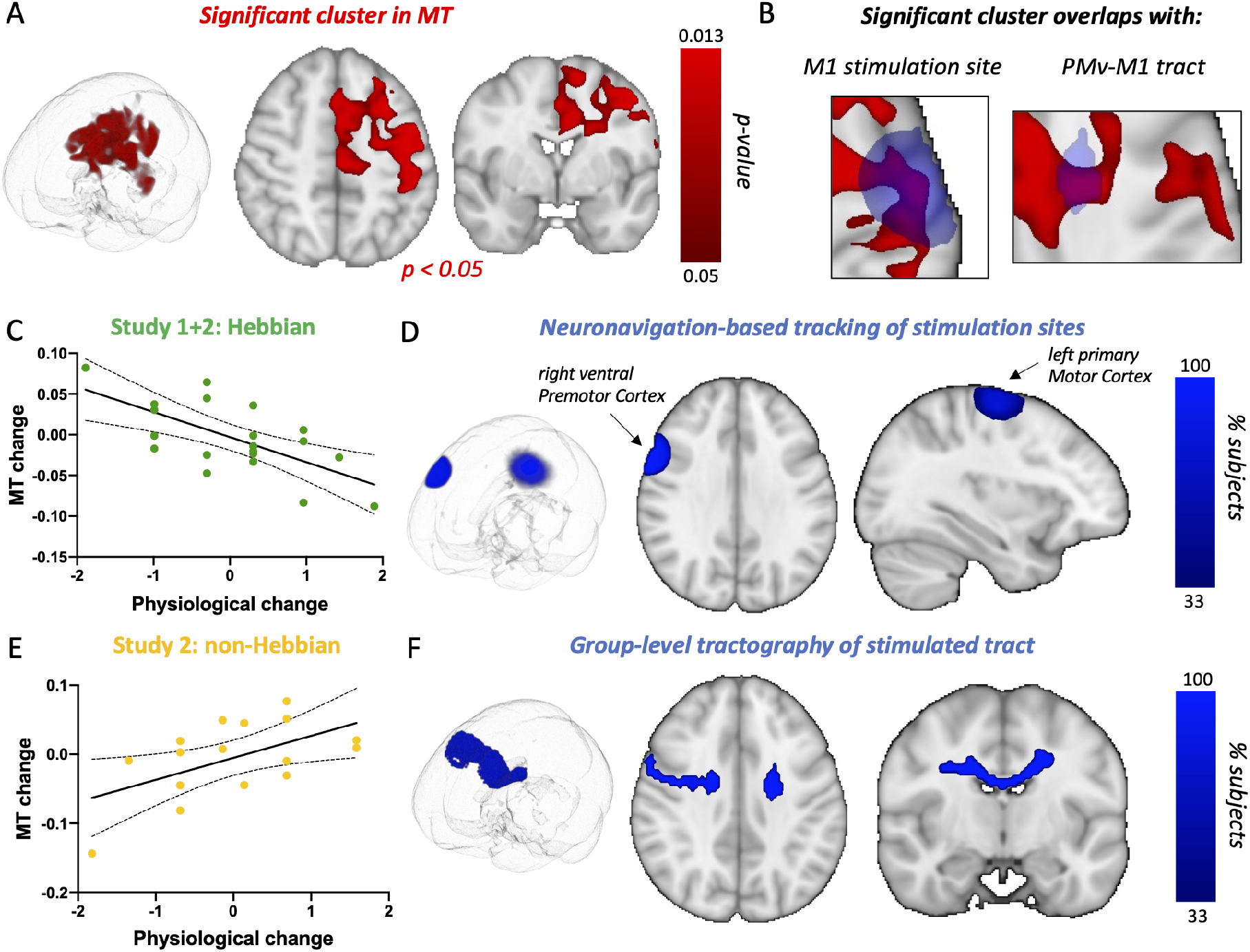
Microstructural Plasticity induced by Hebbian stimulation. A: Results from a whole-brain analysis identify a cluster where changes in MT values correlate with changes in cortical excitability in the Hebbian condition significantly more than they do in the non-Hebbian condition. B: The significant MT cluster identified by the whole-brain analysis (red) overlaps with stimulation sites in the grey matter and with the stimulated fibre tract in the white matter (blue). *C, E*: Scatterplots of data underlying the significant cluster. For the Hebbian condition, participants with greater increases in excitability (more negative physiological change score) show greater increases in MT. Each data point is a single participant; scatterplots (with line of best fit and 95% confidence bands) are presented for post-hoc visualisation of the correlations, rather than for statistical inference. *D, F*: Tracking of stimulation sites via neuronavigation allows to estimate the location of cortical stimulation sites, and to reconstruct the stimulated fiber bundle in white matter.

We investigated the anatomical relationship between Hebbian stimulation and this cluster. We found that in cortical areas, the cluster overlapped with locations of the M1 coil that were recorded during stimulation. We then performed tractography and reconstructed the stimulated white matter bundle connecting the stimulation sites (Fig. S5). The significant cluster overlapped with the reconstructed white matter bundle (Fig. 2B, Fig. 2D, Fig. 2F), further confirming the close relationship between Hebbian stimulation and observed myelin changes.

### Hebbian stimulation induces anatomically-relevant changes in action re-programming

We then assessed whether the Hebbian white matter changes we have observed might play a role in behaviours known to be supported by the white matter fibers being stimulated. In Study 1 and 2, subjects undertook an action reprogramming task (Fig. 3A). Action reprogramming is known to selectively involve the PMv to M1 motor circuit [27], which we further confirmed through a meta-analysis of the action reprogramming task-fMRI literature (Fig. 3B). This is consistent with the observation that not only does PMv have a major projection to M1 [28], which enables it to exert a strong influence over M1 activity [29, 30] but, in addition, PMv receives an especially strong projection from lateral prefrontal cortex [28]. It is also important to note that many PMv projections to M1 terminate on inhibitory interneurons [31, 30]. Thus, in conjunction, PMv’s pattern of anatomical connections ensure that during action reprogramming it can mediate inhibitory influences, originating from executive control processes in prefrontal cortex, over motor processes in M1.

**Figure 3:**
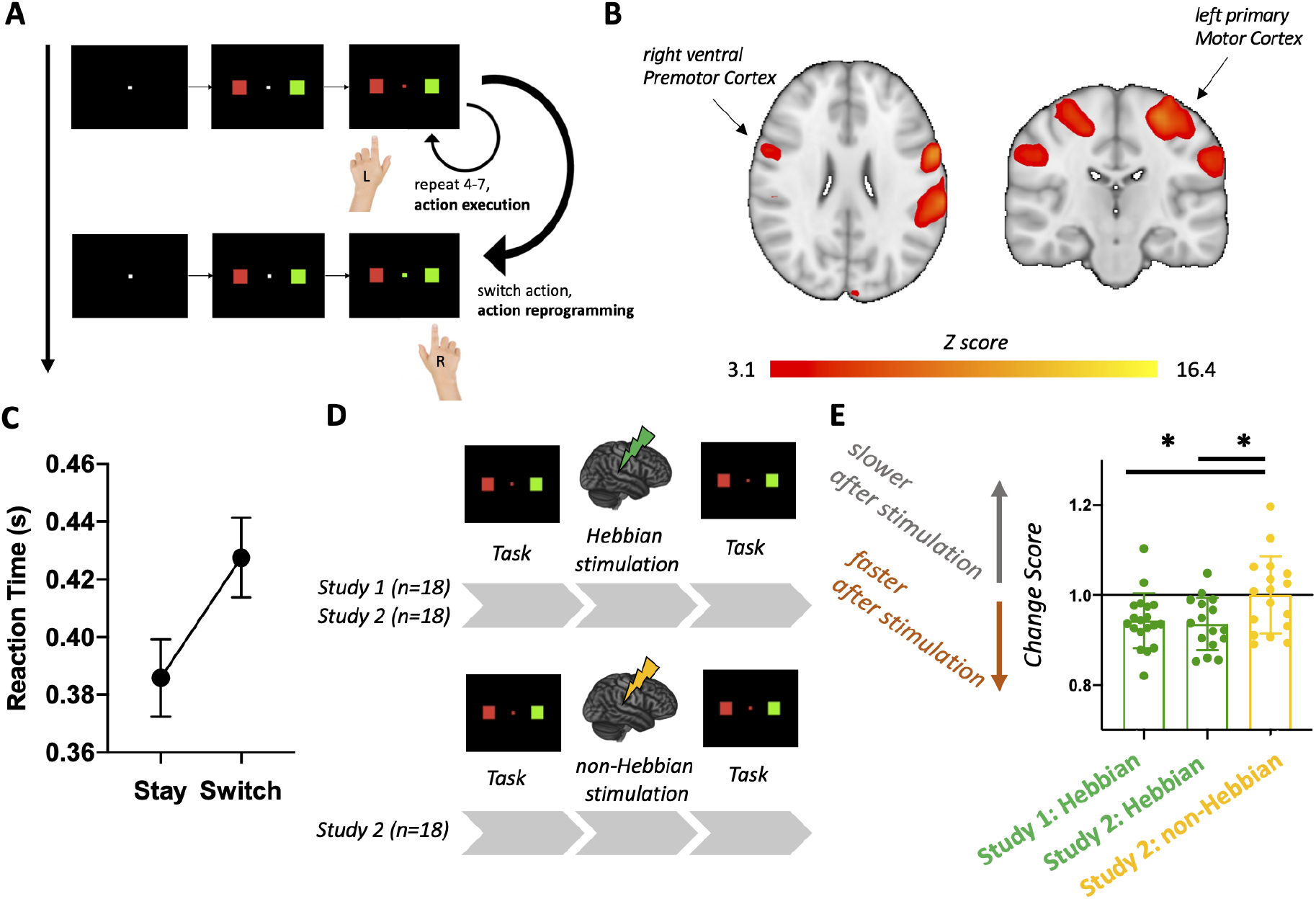
Hebbian stimulation induces anatomically-relevant changes in action reprogramming. *A:* Schematic of the Action Reprogramming Task used, based on [27], probing both action execution (stay trials) and action reprogramming (switch trials). *B*: Premotor-to-motor circuits are involved in action reprogramming, as exemplified by a meta-analysis of action reprogramming task-fMRI studies. *C*: Reaction Times during the task increase in switch trials (when the cue changes) compared to stay trials (while the cue remains the same) in all studies. *D:* Summary of experimental design, testing the effects of Hebbian stimulation on action reprogramming in two cohorts. *E:* Longitudinal effects of Hebbian plasticity induction on action reprogramming behaviour. Each dot in the graph represents the normalized change in switch trial Reaction Time for one subject.

We found that changes in performance on the action reprogramming task differed between the stimulation conditions, specifically on the action reprogramming trials of the task (but not on trials on which participants did not reprogramme actions and simply made the actions that they had pre-prepared; Fig. 3C-E, one-way ANOVA effect of group: F(2, 51)=4.377, p=0.0178). While slower Reaction Times were observed following the control condition, this effect was rescued by the Hebbian plasticity-induction protocol, resulting in relatively improved action reprogramming performance following Hebbian stimulation (post-hoc Study 2:Hebbian vs Study 2:non-Hebbian p = 0.0280; post-hoc post-hoc Study 1:Hebbian vs Study 2:non-Hebbian p = 0.0447). No changes, however, were found when no action reprogramming was required and participants simply made the movements that they had pre-prepared (i.e. stay trials, Fig. S6). Behavioural changes in action reprogramming were present even when covarying for changes in action execution performance during stay trials (one-way ANCOVA effect of group: F(2, 51)=4.373, p=0.018), which further supports a close link between Hebbian stimulation and the observed myelin changes.

### Functional neuroimaging reveals compensatory connectivity changes induced by Hebbian stimulation

Finally, it is possible that Hebbian plasticity may also induce compensatory functional changes [19]. Therefore, we tested whether Hebbian stimulation induces large-scale changes in functional connectivity of the stimulated areas. We found evidence for large-scale compensatory changes in functional connectivity (Fig. 4, peak p_corr_ = 0.001). More specifically, we found that participants with the strongest increase in cortical excitability following Hebbian stimulation also exhibited the strongest decrease in connectivity between stimulated brain areas and non-stimulated visuomotor pathways (Fig. 4A, Fig. 4B), including posterior superior parietal cortex (pSPL) and Area V3A (Fig. S7). This correlation was not present in those receiving non-Hebbian stimulation (Fig. 4C).

**Figure 4:**
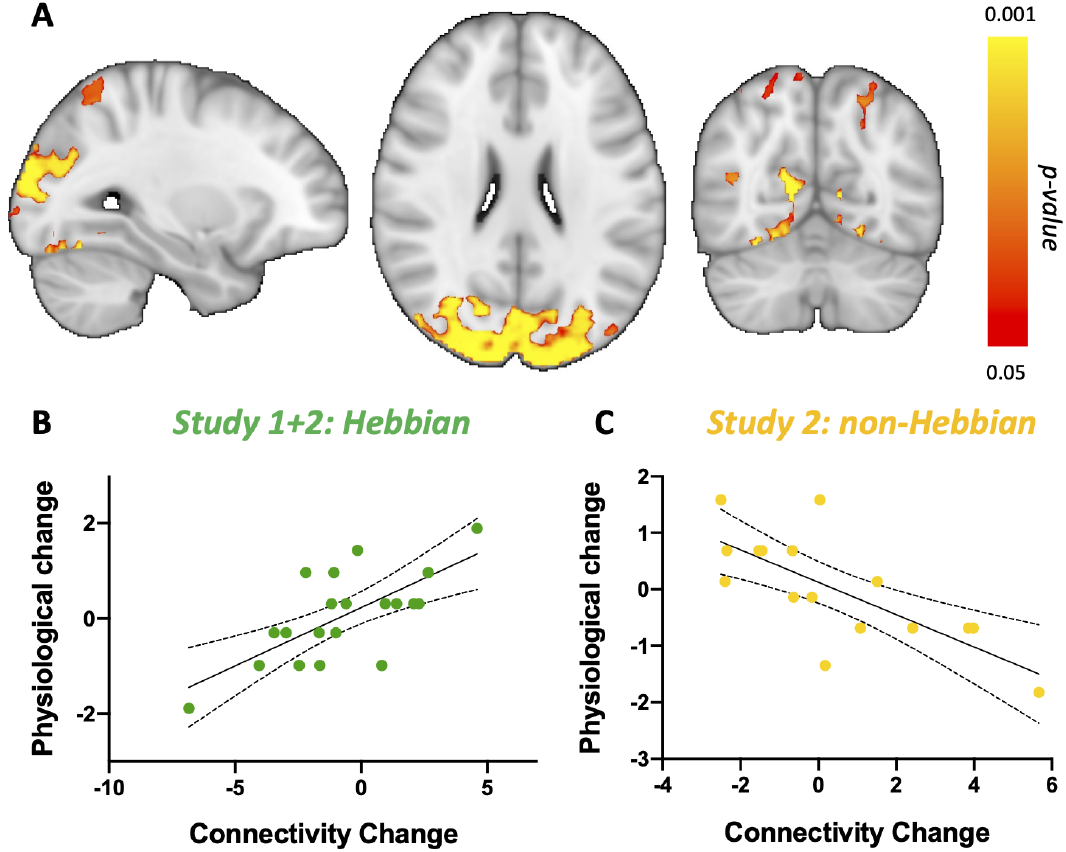
Large-scale compensatory changes in resting-state connectivity induced by Hebbian stimulation. *A:* Results from a whole-brain analysis identify clusters where connectivity changes correlate with changes in cortical excitability in the Hebbian condition significantly more than they do in the non-Hebbian condition. *B, C:* Scatterplots of data underlying the significant cluster. Each data point is a single participant; scatterplots (with line of best fit and 95% confidence bands) are presented for post-hoc visualisation of the correlations, rather than for statistical inference.

## Discussion

Hebb’s Rule provides a rare conceptual link between cellular plasticity (“neurons that fire together, wire together”) and cognition (associative learning), and has thus been central to how we conceive of brain function and learning. While synaptic plasticity has often been assumed to be the cellular basis for Hebbian plasticity [5, 32], here we show that Hebb’s Rule extends beyond synaptic changes. The “neurons that fire together, wire together” principle applies not only to synapses, but also to myelinated long-range connections between neurons, in the white matter.

As Hebbian plasticity requires the detection of coincident neuronal activity, one key implication of our findings is that plasticity in myelinated white matter tracts can be influenced by coincident activity in the areas they connect. While the exact workings of coincidence detection in myelinated axons are unknown, synaptic plasticity and myelin plasticity might rely on the same coincidence detection method. In this scenario, synapses would detect coincident activity and effect changes in myelination, for instance by means of retrograde signalling to the presynaptic axon. An alternative possibility is that myelinating cells themselves might perform coincidence detection. Oligodendrocyte precursors receive direct synaptic input from neurons [33], express NMDA receptors, the same receptors which enable coincidence detection at synapses [34], and can receive inputs from multiple distant but functionally-connected brain areas [35]. Therefore, it is also possible that myelinating cells may directly perform coincidence detection, and that this process may underlie the Hebbian properties of myelin plasticity.

If myelin plasticity is Hebbian, could it also contribute to associative learning? Previous studies have described important contributions of synaptic plasticity to associative learning and memory [5], but also highlighted that impairing synaptic plasticity does not fully abolish associative learning [36, 32]. Our results provide a potential explanation for these mixed findings: additional sites of plasticity may provide pathways by which Hebbian plasticity can still take place without synaptic changes. This is likely to allow associative learning to happen in the absence of canonical synaptic plasticity. Compatible with this hypothesis, recent findings have in fact confirmed that myelin plasticity is necessary for associative learning in a Pavlovian fear conditioning paradigm [16], which may be due to myelin plasticity’s Hebbian properties.

The observation that myelin plasticity is Hebbian also provides new insights regarding what its role in brain function may be. The very existence of myelin plasticity in adulthood has been debated until recently [37, 38], as it is energetically expensive to generate the bulky macro-molecules needed for forming new myelin [39]. While it is now established that myelin changes do happen on the scale of days to weeks [7], it still remains a mystery why the brain might need such a resource-intensive plastic process. Our results highlight that myelin plasticity may have similar computational properties to synaptic plasticity, but unfold over longer timescales. This provides a unique role for myelin plasticity that cannot be fulfilled by synapses alone, and may justify the higher energetic cost needed for the upkeep of myelin plasticity.

Beyond their role in behaviour, another commonality between plasticity in synapses and myelinated axons is that they are both activity-dependent. While in recent years a growing body of research has shown clear evidence that neuronal activity drives myelin changes in rodents [10, 11, 13], our results confirm that white matter plasticity is activity-dependent in humans too. This is noteworthy not only because it proves that key findings from rodent studies can be translated to humans, but also because it opens up the study of activity-dependent myelin plasticity to analyses of interindividual differences. Here, for instance, we show that interindividual differences in cortical excitability explain some variability in the induction of white matter plasticity. This hints that there may be meaningful interindividual variability in activity-driven myelin plasticity, which is unlikely to be detected in genetically and environmentally homogeneous rodent samples [40], but may be accessible in human studies. Moreover, human studies also offer the unique possibility of combining plasticity inductions with *in vivo* functional measurements across the whole brain [41], which, in our case, has allowed us to describe for the first time compensatory functional changes which co-occur with white matter plasticity.

By providing evidence for activity-dependent white matter plasticity in humans, these results bridge two distinct lines of evidence on the topic. Rodent studies have largely focussed on interventional, causal approaches, interrogating the activity-dependent nature of myelin plasticity [10, 11]. By contrast, human studies have focussed on behavioural paradigms and their effects on white matter [8]. Our results bridge these distinct but complementary bodies of research, by showing that activity-dependent white matter plasticity can be induced in humans as well, and studied in conjunction with behaviour. Our observations confirm that translating causal insights from rodents to humans is possible [42, 43, 44], and can bring about new discoveries on the nature and extent of brain plasticity.

Our results further build upon growing evidence that non-invasive imaging can detect subtle microstructural changes. Recent developments in MR physics are allowing researchers to measure quantitative MR parameters with higher reliability than ever before, thus providing new tools to study the relatively subtle changes in myelination that can be experimentally induced in humans. In particular, quantitative markers based on Magnetisation Transfer, such as the one used here, are especially sensitive to the myelin content of a voxel [21], and have been shown to be particularly sensitive to myelin changes in response to behavioural interventions [45]. In summary, non-invasive quantitative markers are not only able to improve our understanding of white matter and myelin plasticity, but may also afford the ability to translate key rodent findings to both healthy and clinical human cohorts [46].

The Hebbian stimulation protocol used here, also known as paired associative Transcranial Magnetic Stimulation (or paTMS), has high translational potential. Most non-invasive brain stimulation protocols have short-lived effects of under an hour [47]. This means that in clinical practice, several stimulation sessions need to be delivered over weeks to observe clinical benefits [48]. By contrast, paTMS induces longer-lasting effects [18], which we show are still present 24 hours after stimulation. This longer time-scale mirrors the longer time-scales of myelin plasticity [7], suggesting that protocols inducing longer-lasting effects, such as the one used here, are particularly promising candidates to induce myelin and white matter plasticity in humans. This hints that brain stimulation protocols aimed at inducing Hebbian plasticity may not only provide much-needed causal insights into basic neuroscience questions, but may also be exploited for clinical use.

Using non-invasive approaches, as we do here, has the important advantage of avoiding confounding effects on glial cells from invasive plasticity inductions [49], but poses limits to our interpretation of the results. There is no 1-to-1 mapping between microstructural MR signals and underlying biology [50]. Therefore, we can infer that there are plastic changes in white matter, and that they are likely driven by myelin, but we cannot distinguish what exact changes in the myelinated axon are causing our observations. For instance, while it is unlikely that *de novo* oligrodendrogenesis would have taken place over 24 hours, we know that remodelling of myelin morphology [22], production of myelin from existing oligodendrocytes [23, 25], or increased concentration of myelin due to shortening of Nodes of Ranvier [24] could have all happened within our experimental time frame, and given rise to our result. A large variety of candidate processes have been proposed to contribute to plasticity of the myelinated axons [7, 14], and while any of them could be driving our observations, our results hint that at least some of them are bound to be Hebbian in nature.

In conclusion, our study combines recent advances in non-invasive brain imaging and brain stimulation to show that Hebb’s Rule extends beyond synapses. While myelin plasticity may provide an additional site of brain plasticity, the same rules may constrain its functions. As our understanding of new forms of brain plasticity develops, we suggest that Hebb’s Rule may be a broader principle than previously thought, constraining multiple plastic processes in the human brain.

## Materials and Methods

### Experimental design of Study 1 and Study 2

All participants underwent three consecutive days of testing (Figure 1A). On the first day, Magnetic Resonance Imaging (MRI) was collected (including myelin markers). On the second day, the participants underwent either Hebbian or non-Hebbian plasticity induction (both achieved through Transcranial Magnetic Stimulation, TMS). On the third day, MRI (including myelin markers) was collected again. Each participant’s sessions were matched to be at the same time of day to control for circadian effects. All participants were self-assessed right-handed and their handedness was further confirmed through the Edinburgh Handedness Inventory [51]. All participants were screened for TMS and MRI safety, received monetary compensation for their participation, and gave their informed consent to participate in this study. All study procedures were reviewed and approved by the local ethics committee at the University of Oxford (Central University Research Ethics Committee (CUREC)), and followed the Declaration of Helsinki.

In study 1, 19 healthy participants (aged 18-32, 9 female) underwent a longitudinal MRI-TMS-MRI paradigm, and all participants underwent the Hebbian plasticity-induction condition.

In study 2, 36 healthy participants (aged 19-30, 22 female) underwent a longitudinal MRI-TMS-MRI paradigm. Participants were randomly assigned either to the Hebbian or the non-Hebbian plasticity induction protocols.

### Hebbian and non-Hebbian Plasticity Induction Protocols

Hebbian and non-Hebbian protocols were both based on paired associative cortio-cortical Tran-scranial Magnetic Stimulation (paTMS), a recently developed stimulation protocol [52, 53, 54, 18, 19] where two cortical regions are repetitively stimulated in a paired fashion at inter-pulse intervals known to induce LTP-like associative synaptic plasticity.

Hebbian (active) and non-Hebbian (control) stimulation protocols both used two DuoMAG MP-Dual TMS monophasic stimulators (DeyMed DuoMag, Rogue Resolutions Ltd.) to deliver paired pulses via two figure-eight coils, one 70mm-diameter coil over primary motor cortex (M1) and one 50mm-diameter coil over ventral premotor cortex (PMv). In the Hebbian condition, the paired pulses were 6ms apart, mimicking the timing of synaptic plasticity inductions used *in vitro*. In the non-Hebbian condition, the pulses were 500 ms apart, which is long enough to avoid physiological interactions between the two pulses which may take place at shorter intervals [55]. Moreover, using a 500 ms interval has been shown not to have behavioural and physiological effects in previous studies [19]. All other parameters were the same across protocols: both protocols consisted of 90 paired TMS pulses, delivered at 0.1 Hz over a 15 minute period, without interruptions. For both protocols, the M1 coil was set at a SI_1mV_ intensity, whereas the ventral premotor cortex coil was set at a 110% resting Motor Threshold intensity (rMT). SI_1mV_ was determined as the intensity giving reliable and stable 1 mV Motor-Evoked Potentials (MEPs) at rest over 10 pulses. rMT was determined as the intensity at which 5 out of 10 pulses gave no MEP response greater than 0.05 mV.

Both protocols were performed at rest, with the participant resting their hands on a pillow and watching a series of still images on a computer screen. In summary, each area, PMv and M1, was stimulated in an identical manner in the two protocols; each was stimulated the same number of times at the same intensity and frequency and for the same duration as in the other protocol, but the relative timing of stimulation meant that spike-timing-dependent plasticity could only occur in one protocol.

Subjects were blind to their experimental condition throughout the experiment. Experimenters were also blind to the experimental condition prior to stimulation; however, the subtle difference in stimulation timing between Hebbian stimulation and non-Hebbian stimulation made it impossible to achieve full blinding once the stimulation had started taking place. At the end of the experiment, participants were administered a discomfort questionnaire [74] and a questionnaire aimed at assessing blinding of the experiment (both available here: https://open.win.ox.ac.uk/pages/alazari/hebbian-white-matter-plasticity/). The scores from the blinding questionnaire were used to calculate an overall Bang’s Blinding Index for the experiment [76].

### Neuronavigation

All stimulation was delivered using continuous tracking of coil location with respect to subject neuroanatomy (i.e. neuronavigation). This was achieved through a Polaris camera and the Brainsight software (Rogue Resolutions, Inc.), and used the participants T1-weighted (T1w) structural MRI scan. The participant was tracked via a headband with reflective spheres attached to it; the coils were tracked with coil trackers that were re-calibrated at the beginning of each testing day. Online neuronavigation ensured that all stimulation sites were within 3 mm of target location, as described in previous publications [18].

Coil location was also recorded and analysed offline. An automated Brainsight tool was used to find the closest brain voxel to the sampled stimulation site. The coordinates for this voxel were then transformed to standard space to allow overlaying of stimulation sites from different participants. At this stage, a total of 42 stimulation locations were included, as 4 participants’ stimulation locations failed to save due to software fault (2 in active-only study, 1 in active randomised and 1 in control randomised), and 5 participant’s stimulation locations could not be automatically determined with Brainsight (2 in active-only, 2 in active randomised, 1 in control randomised). Because the magnetic field may reach 30% of its peak level throughout a region with a diameter of 4 cm [73], spheres of 4 cm diameter were created around the sample stimulation location to provide a conservative estimate of the spatial specificity achieved by TMS. These spheres were then overlaid upon each other. All stimulation sites were within 3 mm of target location, as described in previous publications [18].

### Cortical Physiology

As a measure of cortical excitability, we determined the Stimulator Intensity giving reliable and stable 1 mV Motor-Evoked Potentials in the First Dorsal Interosseus muscle of the right hand (‘SI_1mV_’) [56]. The SI_1mV_ value was determined at rest and based on 10 TMS pulses. This measure was collected before Hebbian stimulation on day 2 and before MRI scanning on day 3, taking care that sessions were matched to be at the same time of day to control for circadian effects. The SI_1mV_ value was collected in an exploratory manner in the last 7 participants of study 1, and in all participants of study 2 to confirm the presence of longitudinal effects.

### Magnetic Resonance Imaging of Myelin

Participants underwent Magnetic Resonance Imaging (MRI) sessions 24 hours before and 24 hours after the plasticity induction protocol. MRI data were collected with a 3.0-T Prisma Magnetom Siemens scanner, software version VE11C (Siemens Medical Systems, Erlangen, Germany). T1-weighted structural imaging (T1w), Diffusion-Weighted Imaging (DWI) and Multi-Parameter Mapping (MPM) sequences were collected.

The T1w sequence (TR = 1900 ms, TE = 3.96 ms, resolution = 1 mm isotropic, GRAPPA = 2) had a large Field of View (FOV = 256 mm^3^)to allow for the nose and intertragic notches of the ears to be included in the image to facilitate later neuronavigation of the TMS coil to the target position.

Diffusion-weighted Echo-planar imaging (EPI) scans (TR = 3070 ms, TE = 85.00 ms, FOV = 204 mm^3^, resolution =1.5 mm isotropic, multiband factor of 4) were collected for two b-values (500 and 2000 s/mm^2^), over 281 directions. An additional 23 volumes were acquired at b=0, 15 in anterior-posterior (AP) phase-encoding direction and 8 in the posterior-anterior (PA) phase-encoding direction.

The MPM protocol [57] included three multi-echo 3D FLASH (fast low-angle shot) scans with varying acquisition parameters, one RF transmit field map (B1+map) and one static magnetic (B0) field map scan, for a total acquisition time of roughly 22 minutes. To correct for inter-scan motion, positionspecific receive coil sensitivity field maps, matched in FOV to the MPM scans, were calculated and corrected for [58]. The three 3D FLASH scans were designed to be predominantly T1-, PD-, or MT-weighted by changing the flip angle and the presence of a pre-pulse: 8 echoes were predominantly Proton Density-weighted (TR = 25 ms; flip angle = 6 degrees; TE = 2.3-18.4 ms), 8 echoes were predominantly T1-weighted (TR = 25 ms; flip angle = 21 degrees; TE = 2.3-18.4 ms) and 6 echoes were predominantly Magnetisation Transfer-weighted (MTw, TR = 25ms; flip angle = 21 degrees; TE = 2.3-13.8 ms). For MTw scans, excitation was preceded by off-resonance Gaussian MT pulse of 4 ms duration, nominal flip angle, 2 kHz frequency offset from water resonance. All FLASH scans had 1 mm isotropic resolution, field of view (FOV) of 256 x 224 x 176 mm^3^, and GRAPPA factor of 2×2. The B1 map was acquired through an EPI-based sequence featuring spin and stimulated echoes (SE and STE) with 11 nominal flip angles, FOV of 192 x 192 x 256 mm^3^ and TR of 500 ms. The TE was 37.06 ms, and the mixing time was 33.8 ms. The B0 map was acquired to correct the B1+ map for distortions due to off-resonance effects. The B0 map sequence had a TR of 1020.0 ms, first TE of 10 ms, second TE of 12.46 ms, field of view (FOV) of 192 x 192 x 256 mm^3^ and read-out bandwidth of 260 Hz/pixel.

MRI scan pre-processing, analysis and statistical comparisons were performed using FMRIB Software Library (FSL, v6.0) [59], except for the MPM quantitative map estimation step which was carried out using the hMRI toolbox implemented in Matlab-based SPM, as described in [60]. All T1w images were preprocessed through a standard FreeSurfer-based pipeline [61, 62] to correct for bias field and achieve ACPC alignment (for use in Neuronavigation). For longitudinal analyses of MRI, a midpoint T1w space was derived as done in previous studies [8].

Custom pipelines based on existing FSL tools were developed to preprocess diffusion and Magnetisation Transfer saturation (MT) data (code available here: https://open.win.ox.ac.uk/pages/alazari/hebbian-white-matter-plasticity/). For diffusion, the *topup* tool was run on average images of AP b0 volumes and PA b0 volumes. The resulting susceptibility-induced off-resonance field was then used as an input for the eddy tool [63], which was run with options optimised for multiband diffusion data to correct for eddy currents and subject movement.

Magnetisation Transfer saturation (MT) quantitative maps were estimated through the hMRI toolbox [60]. MPM volumes were then registered to Montreal Neurological Institute (MNI) space by combining the registration between MPM volumes and midpoint T1w images with the registration between the midpoint T1w space and the MNI template; these volumes were then smoothed with a Gaussian kernel of 3 mm. At this stage, 1 participant was excluded as their MPM scans were heavily corrupted due to movement artefacts (study 1); 1 participant was excluded due to lower quality signal in the MPM scans, which resulted in poor registration to template (study 2, control condition); 1 participant was excluded due to a slight callosal abnormality preventing registration to template (study 2, active condition).

### Magnetic Resonance Imaging of Resting-State Connectivity

The rs-fMRI Echo-planar imaging (EPI) sequence (TR = 750 ms; TE = 29.00 ms, resolution = 2 mm isotropic, FOV = 208 mm^3^) employed fat saturation-based fat-suppression, had a multiband factor 6 and used GRAPPA with acceleration factor 2. Participants were asked to keep their eyes open and let their mind wander during this sequence. The screen was kept black for the duration of this scan, and an eye tracker was used to ensure the participant was awake. rs-fMRI data was preprocessed with high pass filter cutoff of 100 s, MCFLIRT to correct for motion, smoothing at sigma of 3 mm, BBR registration, and fieldmap-based B0 unwarping. Single session Multivariate Exploratory Linear Optimized Decomposition into Independent Components (MELODIC)-based Independent Component Analysis was used to extract components at the single subject level. Components were classified as noise or signal manually for the first few subjects, and the labels were then used to build a FIX-based classifier to denoise the data [64]. Finally, dual regression [65] was used to estimate connectivity maps from the stimulated ROIs.

### Statistical Inference for Magnetic Resonance Imaging data

Group-level analyses of MT and rs-fMRI maps were conducted through nonparametric permutation inference in the *randomise* tool [66], controlling for the family-wise error rate. Change maps were calculated for each subject by subtracting the day 3 map from the day 1 map. Analyses were run in MNI space, with 10,000 permutations and Threshold-Free Cluster Enhanced [67]. Membership of different studies (Study 1 and Study 2) was encoded as a covariate, to allow for contrasts to test whether effects were present in each study. Changes in the SI_1mV_ metric of cortical physiology (see above) were used as the regressor of interest, explicitly testing for interactions between Hebbian and non-Hebbian conditions. In a separate model, group-level differences between Hebbian and non-Hebbian conditions were also tested through an unpaired t-test.

### Reconstruction of Stimulated Fiber Bundles

Using Diffusion-Weighted Imaging (DWI) data, we reconstructed the white matter fiber bundles stimulated in our plasticity induction protocols. White matter bundles connecting the stimulated cortical sites were estimated using multi-fibre probabilistic diffusion tractography through Prob-trackx [68]. Regions of interest (ROI) in the cortex were based on the neuronavigation-derived sites for each participant, as described above. As the motor hotspot does not always overlap with the postcentral gyral fold, and a larger coil was used for M1 compared to PMv, the motor hotspot ROI was enlarged to a 3cm radius to improve the output tract quality. Tractography was run in native DWI space, with outputs in Montreal Neurological Institute (MNI) space to enable pooling of results across all subjects. Individual-level maps of streamline densities were thresholded at 1% of the number of total valid streamlines per subject, binarised, and then overlaid.

### Action Reprogramming Task

Participants completed the Action Reprogramming task before and immediately after plasticity induction. The Action Reprogramming task aimed to probe action execution and action reprogramming [69, 27]. Cues consisted of a central square (either red or green) with two ‘flanker’ squares (one red and one green). Participants were instructed to press the button on the side where the flanker colour matched the colour of the central square. The flankers kept switching sides at random, whereas the central square was the same colour for 3-7 trials at the time (Figure 3). This way, participants simply had to execute a movement when the central cue colour stayed the same (‘stay trials’), but they had to inhibit the movement and carry out a different one in the trials where the central cue colour had switched (‘switch trials’). Each participant underwent a session of 112 switch trials and corresponding stay trials (for a total of 678 trials). Participants were told to be as fast and accurate as they could. They received detailed task instructions in paper format at the beginning of each session; in addition, the instructions were reiterated in a computer-based fashion at the beginning of the baseline task (code available here: https://open.win.ox.ac.uk/pages/alazari/hebbian-white-matter-plasticity/). Before the baseline task, they undertook roughly 100 trials to make sure first that they understood the rules of the task, and that they had habituated to the task. Three participants (1 in study 2, control condition; 2 in study 2, active condition) had recently performed the same action reprogramming task extensively as part of a separate experiment, and were thus excluded from behavioural analyses to avoid the possibility of training and/or carry-over effects.

### Action Reprogramming Meta-Analysis

A meta-analysis of task-based neuroimaging studies involving action reprogramming was run using the NeuroQuery tool [77]. NeuroQuery performs multivariate prediction-based metaanalyses using text-based search terms and produces meta-analytic activation maps that refer to the concept of interest. As the previous literature consistently refers to our concept of interest as ‘action reprogramming’ [27], we used this as the search term for our meta-analysis.

### Statistical inference for Cortical Physiology and Action Reprogramming data

Analyses of cortical physiology (with SI_1mV_ as the key variable of interest) and Action Reprogramming behaviour (with Reaction Times as the key variables of interest) were run in GraphPad Prism (GraphPad Software, La Jolla, California, US) with the exception of the ANCOVA analysis of Reaction Times which was run in SPSS (SPSS Statistics, IBM Corp.). Longitudinal analyses across all groups were run as one-way ANOVAs of longitudinal change scores, with Dunn’s multiple comparison tests as post-hoc tests. Longitudinal analyses across all groups which aimed to covary for additional factors were run as ANCOVAs. Alpha level for statistical significance was set at 0.05 and all Confidence Intervals (CIs) were set at 95% confidence.

## Acknowledgments

We are grateful to Tim Behrens, Jason Lerch, Antoine Cherix, Laia Serratosa Capdevila, Ru-airi Roberts, Alex S. Bates, Yajing Xu and Claire Bratley for their input on the manuscript. We would like to thank Juliet Semple, Nicola Aikin and Sebastian Rieger for their technical support and help with scanning participants. We acknowledge the IT-related support provided by Matthew Webster, David Flitney and Duncan Mortimer throughout the project. We thank Stuart Clare, Cassandra Gould Van Praag, and Sebastian Rieger for facilitating the sharing of materials as part of this study. This work was supported by a PhD Studentship awarded to AL from the Wellcome Trust (109062/Z/15/Z) and by a Principal Research Fellowship from the Wellcome Trust to HJB (110027/Z/15/Z). The project was supported by the NIHR Oxford Health Biomedical Research Centre and the NIHR Oxford Biomedical Research Centre. The Wellcome Centre for Integrative Neuroimaging is supported by core funding from the Wellcome Trust [203139/Z/16/Z]. This research was funded in whole, or in part, by the Wellcome Trust [Grant numbers 109062/Z/15/Z and 110027/Z/15/Z]. For the purpose of Open Access, the author has applied a CC BY public copyright licence to any Author Accepted Manuscript version arising from this submission.

## Supplementary Results

**Supplementary Figure 1:**
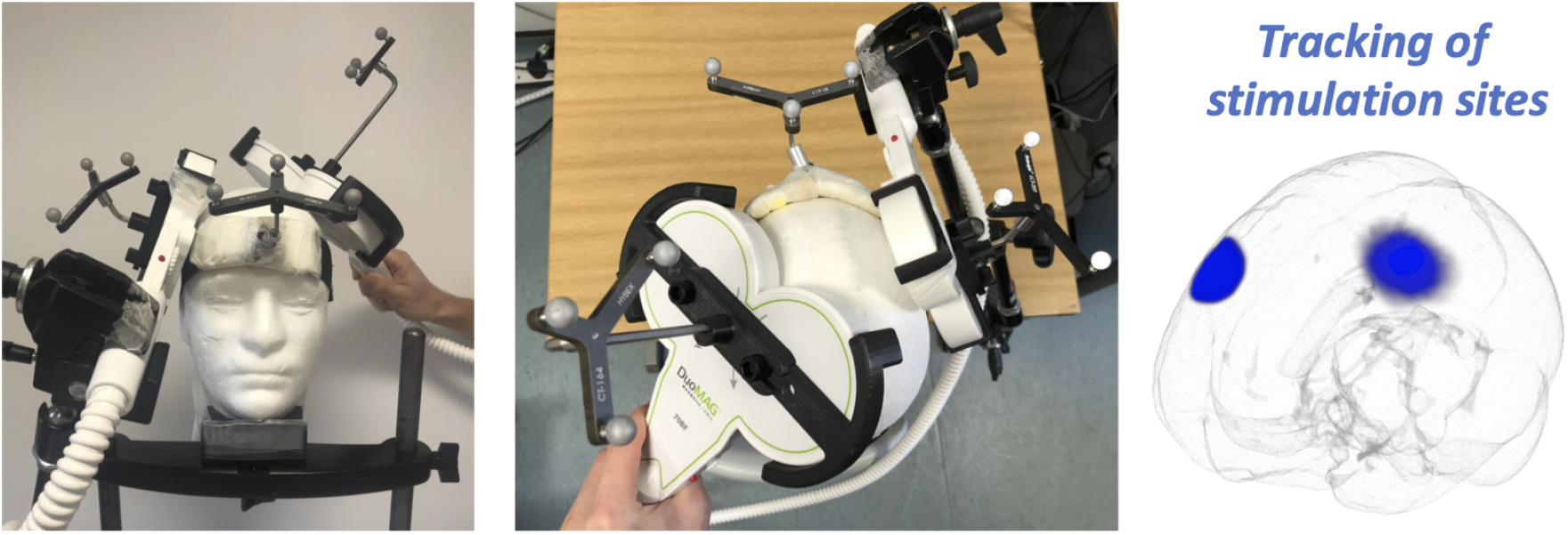
Neuronavigation set-up. *Left:* All stimulation was delivered using continuous tracking of coil location with respect to subject neuroanatomy (i.e. neuronavigation), which was achieved through reflective sphere attached to headbands and coil holders. Online neuronavigation ensured that all stimulation sites were within 3 mm of target location. *Right:* Coil location was also recorded and used for further analyses offline. Here, the stimulation location for all subjects are overlaid in a single 3D image.

**Supplementary Figure 2:**
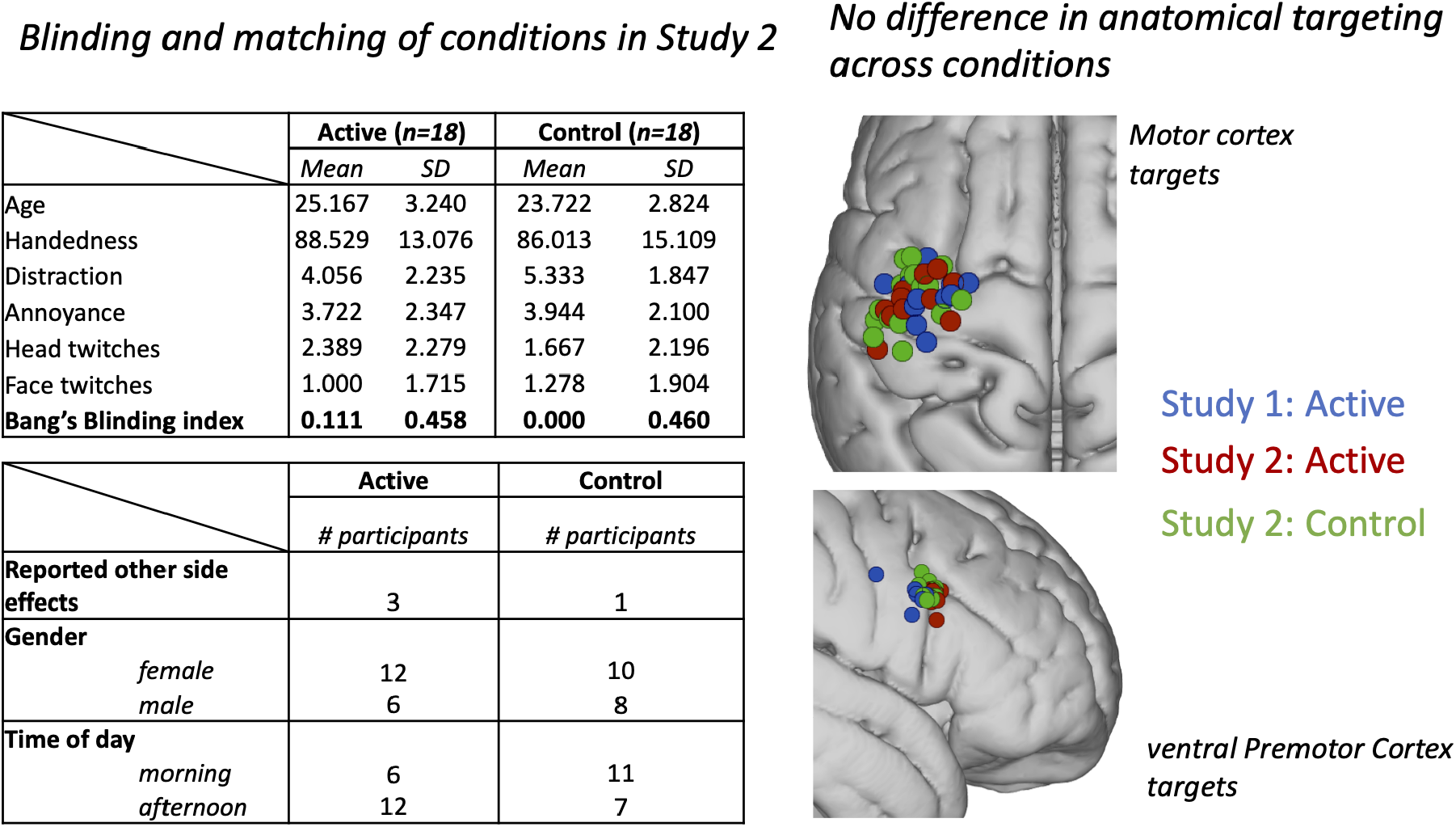
Randomisation and Blinding. *Left:* In Study 2, participants in the Hebbian (active) and Non-Hebbian (control) groups were well matched for demographic variables such as age and gender. Furthermore, the two stimulation protocols did not lead to different experiences of stimulation side-effects, and blinding was successful in both stimulation conditions. *Right:* Offline analysis of neuronavigation target locations shows similarity in the anatomical targeting of stimulation across participants in Study 1 and Study 2, independently of the assigned condition. PMv stimulation was targeted in an anterior position on the boundary between ventral area 6 and area 44 [70, 71] adjacent to the inferior precentral sulcus.

**Supplementary Figure 3:**
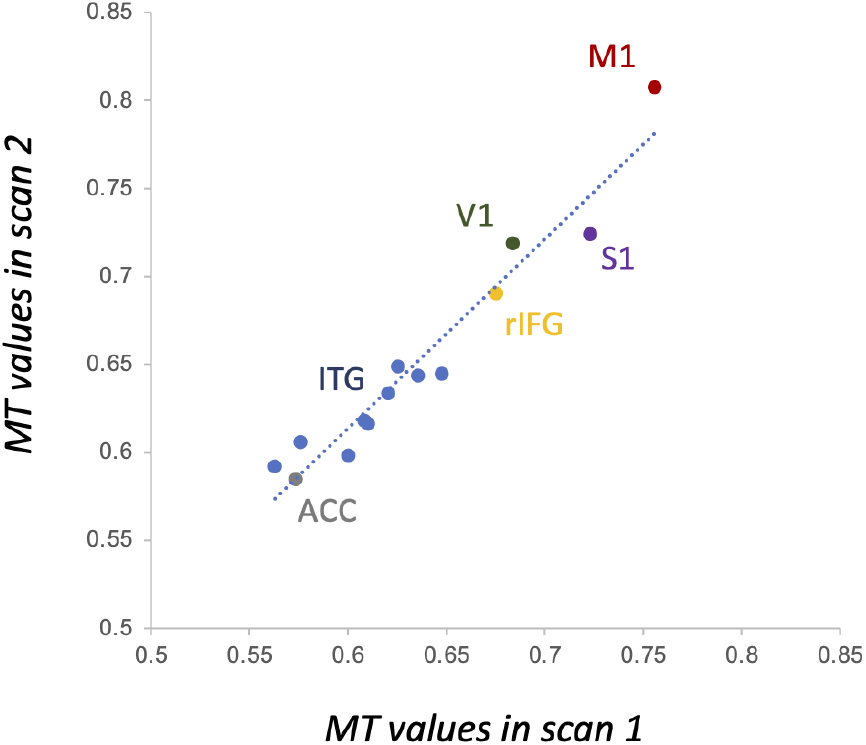
MT has high test-retest reliability on the same scanner across different days. One subject (author A.L.) underwent MPM scanning on the same scanner on different weekdays. Values from MT maps across a range of ROIs have high test-retest reliability within the same scanner.

**Supplementary Figure 4:**
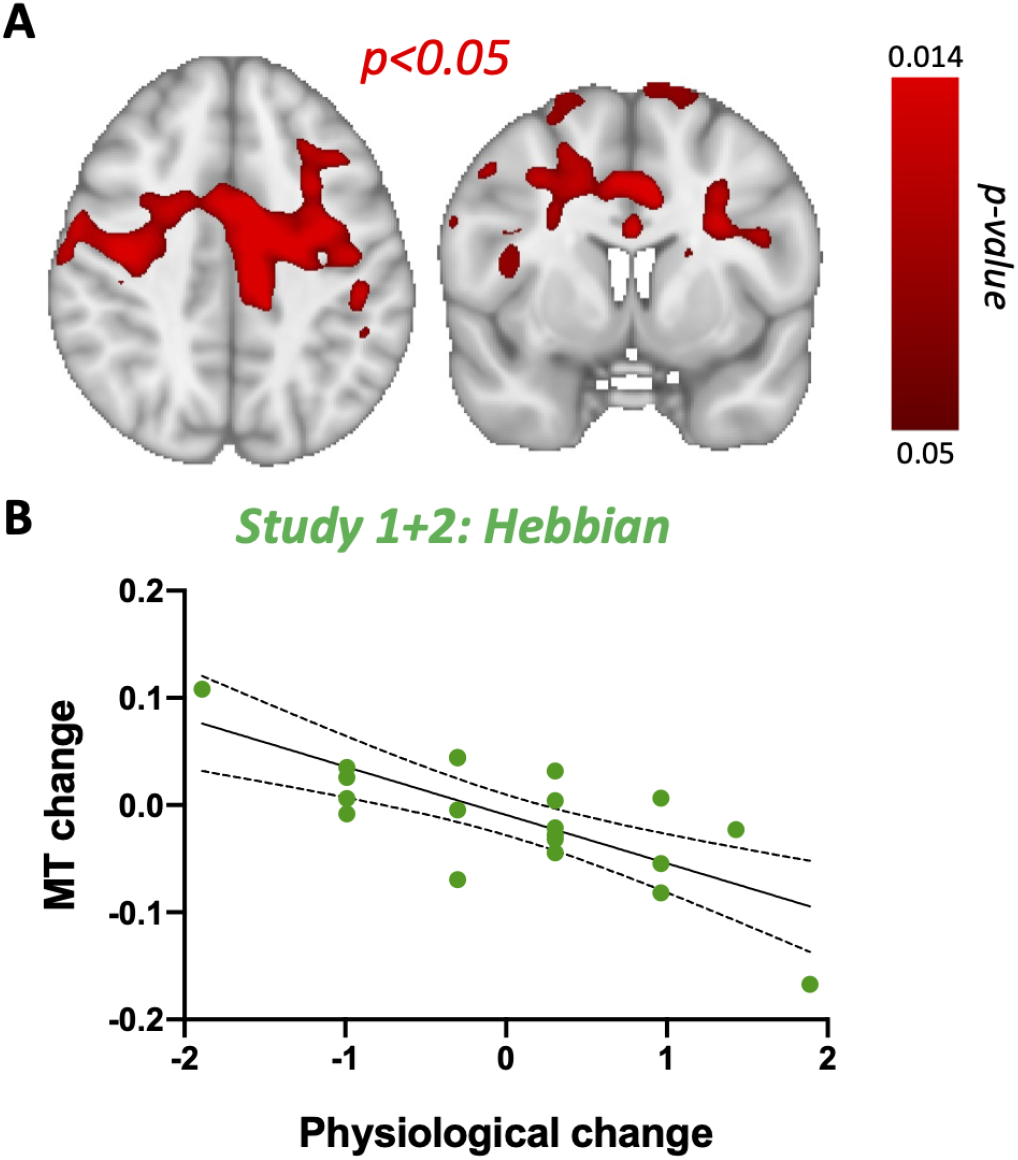
Microstructural results in Hebbian subjects only. To further confirm the correlation between physiological change and MT change, we ran a voxelwise analysis on MT using physiological change as a regressor in Hebbian subjects only (across Study 1 and Study 2). This generates a similar cluster of significant correlation, extending across both hemispheres (A), where greater increases in excitability (more negative physiological change score) are associated with greater increases in MT. This confirms that physiological change is correlated with MT change in white matter, even when considering the Hebbian group alone without contrasting this correlation with the one in the control group.

**Supplementary Figure 5:**
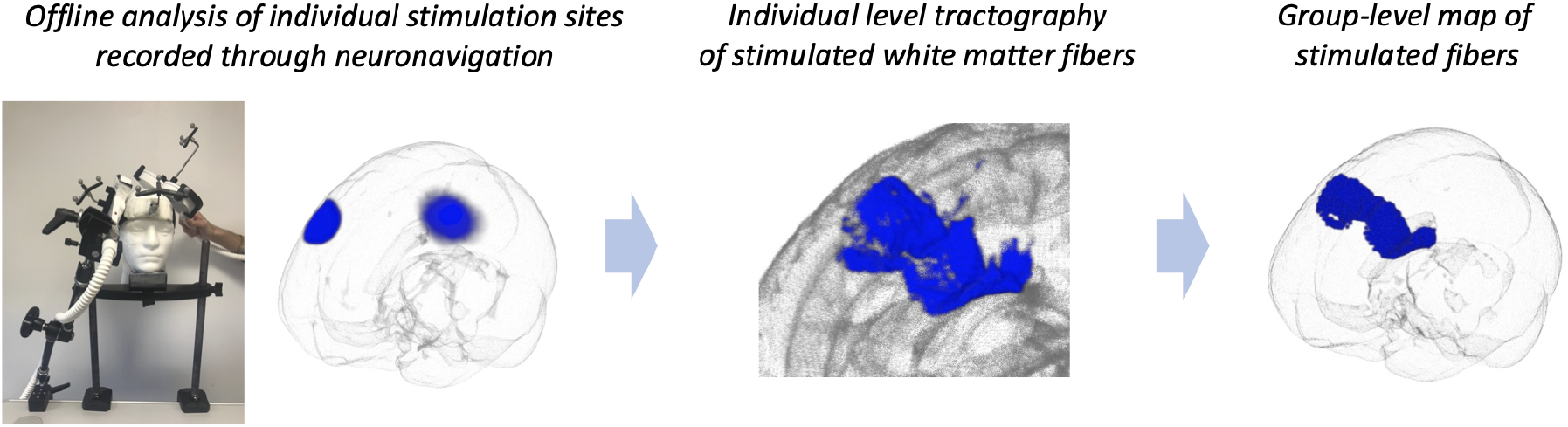
Tractography of stimulated white matter fibres. Left: Neuronavigation allows recording of exact sitmulation locations for each subject. *Centre:* Using stimulation locations for each individual, we estimate individual-level white matter fibers stimulated in our paradigm. *Right*: The group-level map of tract overlap across subjects is shown in blue.

**Supplementary Figure 6:**
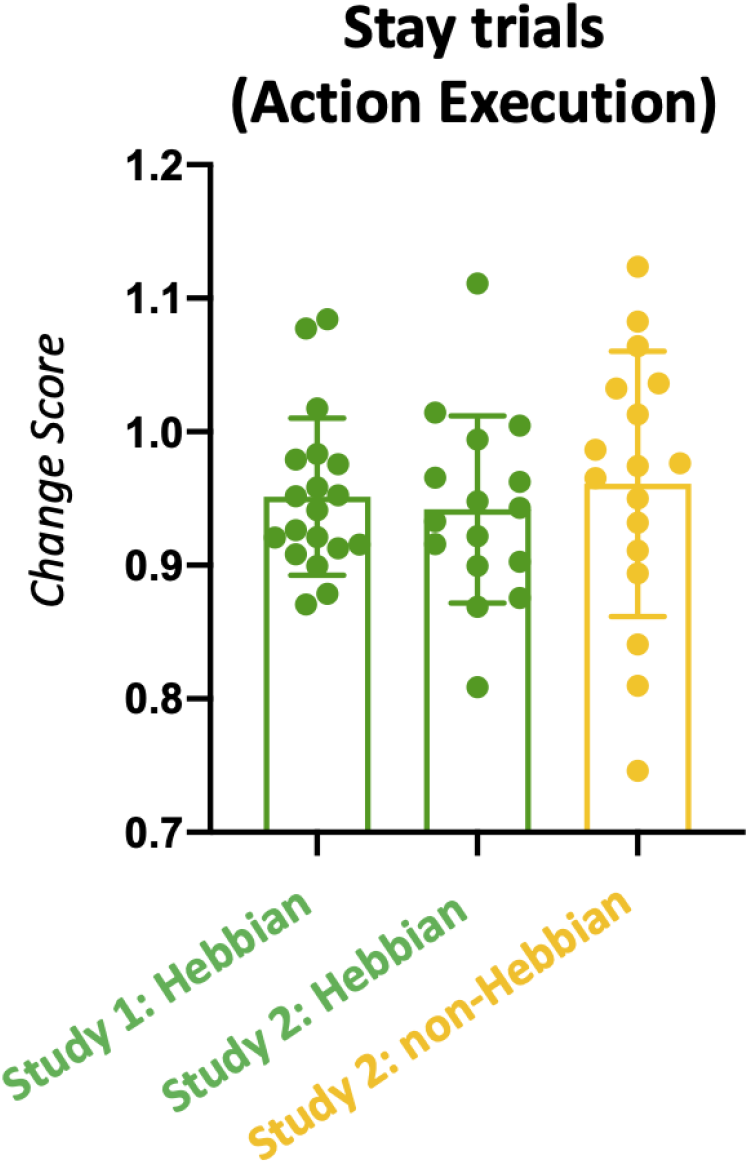
Behavioural effects of Hebbian stimulation do not extend to action execution. Each dot in the graph represents the change in stay trial Reaction Time for one subject. When considering stay trials (action execution), no significant difference was found between groups (one-way ANOVA effect of group: F(2, 51)=1.100, p=0.5769).

**Supplementary Figure 7:**
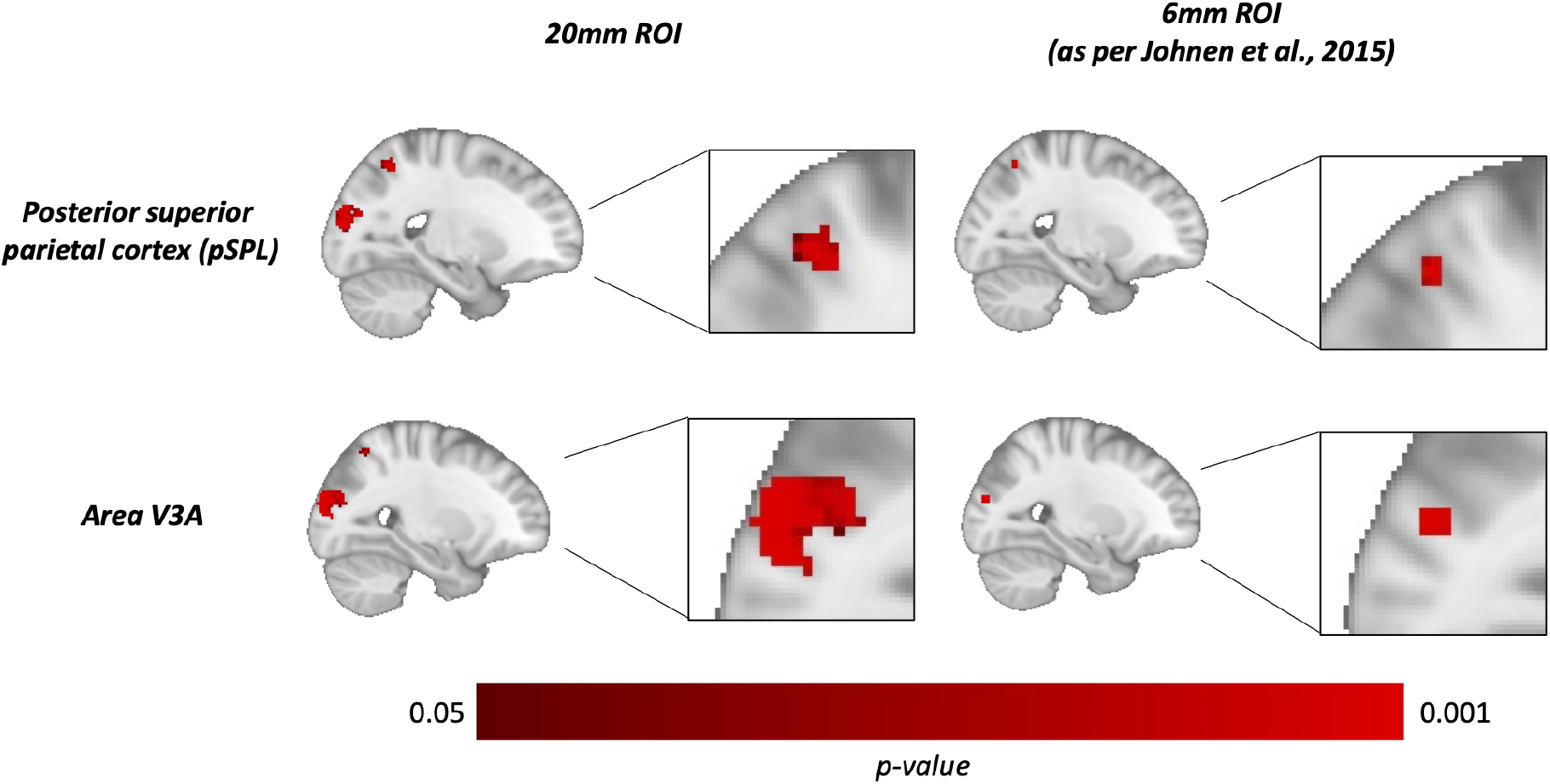
Large-scale compensatory changes in resting-state connectivity, explored through ROI analyses.

## Notes

https://open.win.ox.ac.uk/pages/alazari/hebbian-white-matter-plasticity/

### Competing Interest Statement

The authors have declared no competing interest.

